# Dataset on the mass spectrometry-based proteomic profiling of mouse embryonic fibroblasts from a wild type and DYT-TOR1A mouse model of dystonia, basally and during stress

**DOI:** 10.1101/2021.07.09.451632

**Authors:** Kunal Shroff, Zachary F. Caffall, Erik J. Soderblom, Greg Waitt, Tricia Ho, Nicole Calakos

## Abstract

Here, we present quantitative subcellular compartment-specific proteomic data from wildtype and DYT-TOR1A heterozygous mouse embryonic fibroblasts (MEFs) basally and following thapsigargin treatment [1]. In this experiment, we generated MEFs from wild type and a heterozygous DYT-TOR1A mouse model of dystonia. Subsequently, these MEF cultures were treated with either 1 μM thapsigargin or dimethylsulfoxide vehicle for six hours. Following treatment, the cells were fractionated into nuclear and cytosolic fractions. Liquid chromatography, tandem mass spectrometry (LC/MS/MS)-based proteomic profiling identified 65,056 unique peptides and 4801 unique proteins across all samples. The data presented here provide subcellular compartment-specific proteomic information within a dystonia model system both basally and under cellular stress. These data can inform future experiments focused on studying the function of TorsinA, the protein encoded by TOR1A, and its potential role in nucleocytoplasmic transport and proteostasis. In addition, the information in this article can also inform future mechanistic studies investigating the relationship between DYT-TOR1A dystonia and the cellular stress response to advance understanding of the pathogenesis of dystonia.

## Specifications Table

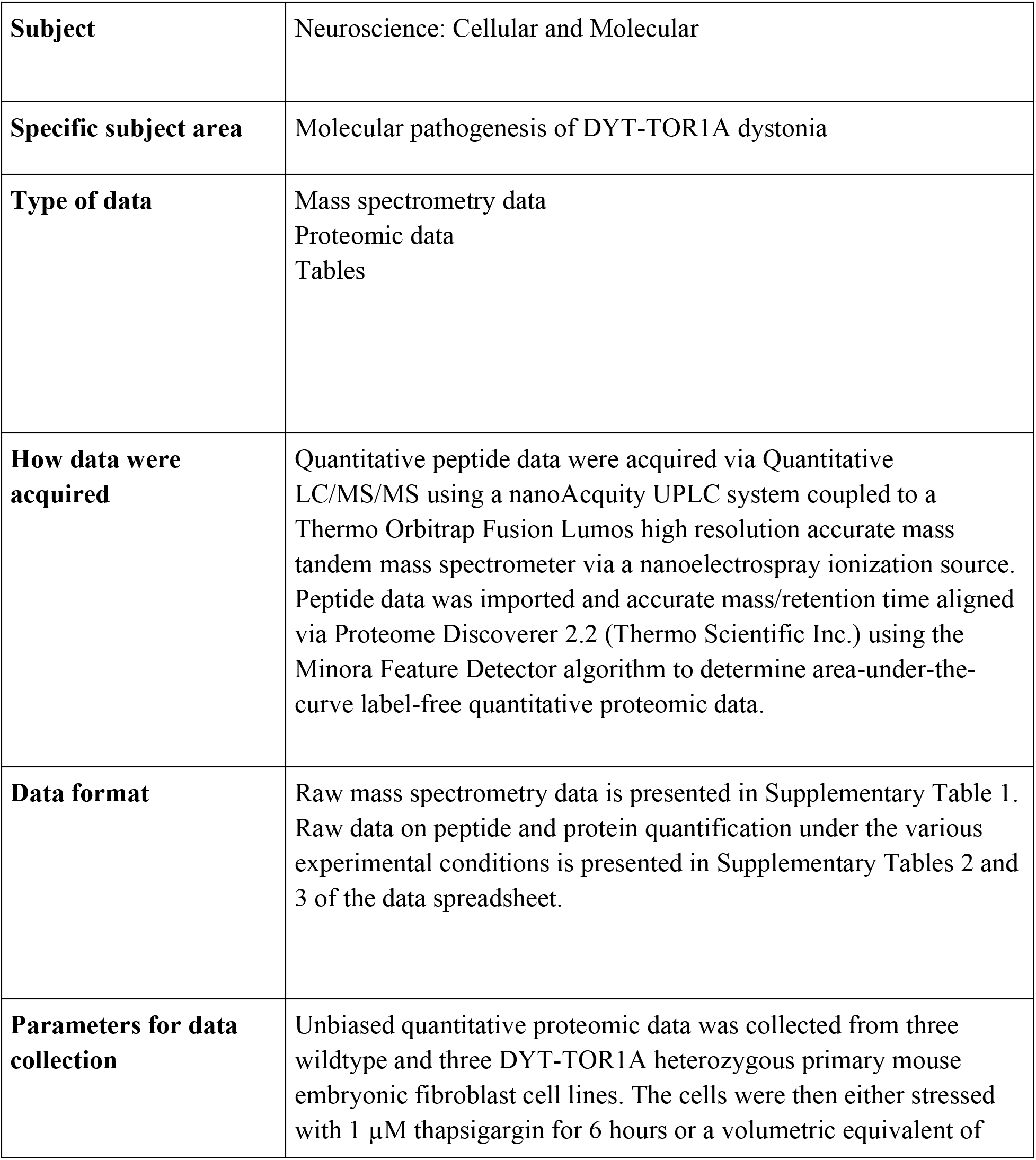
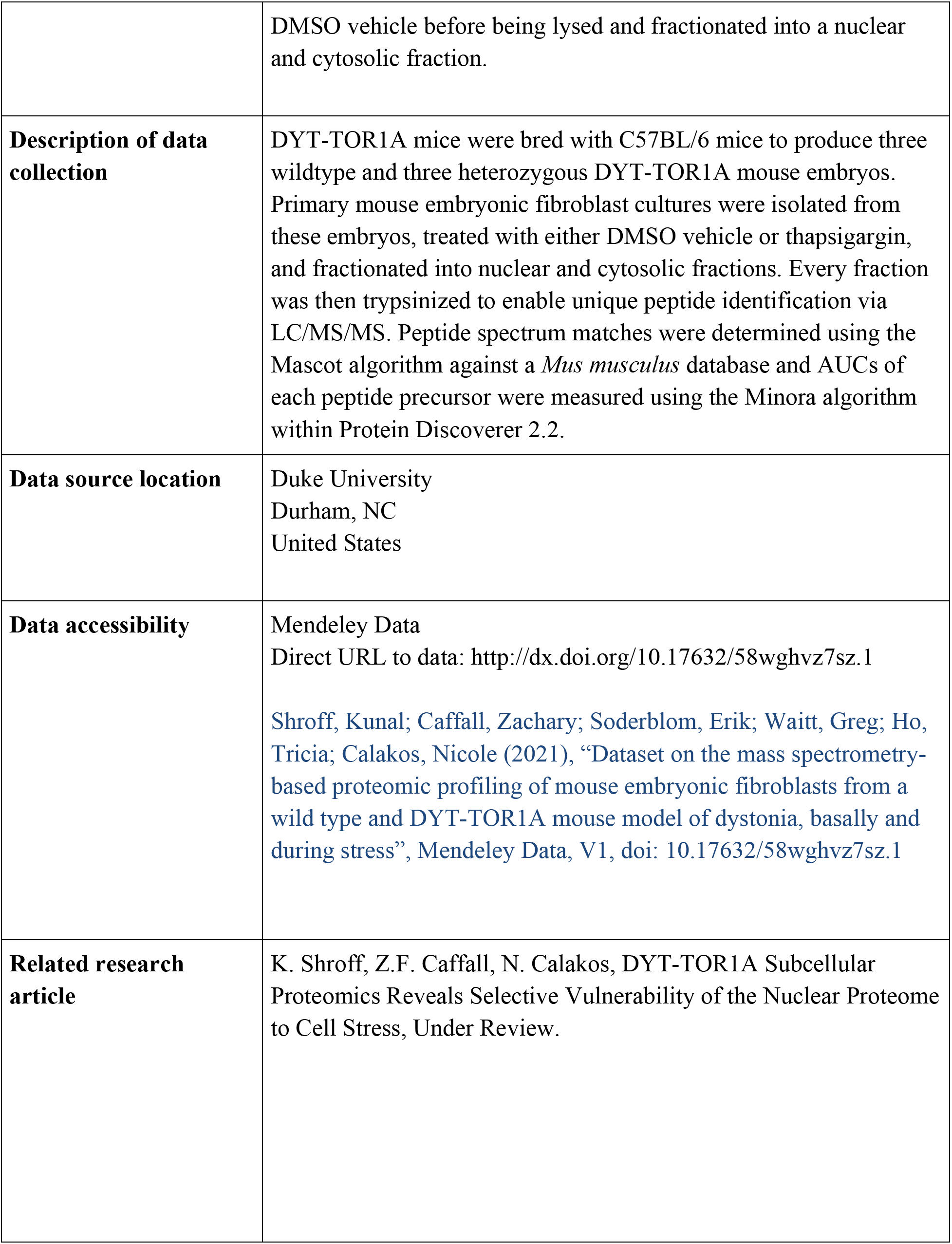

## Value of the Data

- Our data provide information about subcellular compartment-specific proteome changes in DYT-TOR1A dystonia cell models. Because the TorsinA protein traffics between two subcellular compartments, an analysis separating the cytosolic and nuclear compartments is useful to identify disruptions that are specific to a compartment and could be missed by evaluating unfractionated cells or tissue.
- These data can be beneficial to researchers interested in: cell biology of the AAA^+^-ATPase TorsinA, biology of cell stress responses, and understanding how the DYT-TOR1A mutation alters the cellular proteome to gain insights to dystonia pathogenesis.
- These data can be further used to develop experiments aimed at understanding the function of TorsinA and investigating its potential role in regulating nucleocytoplasmic transport and/or proteostasis.
- These data can be used to identify novel proteins that may be mediators of TOR1A’s pathogenic effects particularly when expressed in combination with exogenous cellular stress.

## Data Description

We generated quantitative proteomic profiles of the nuclear and cytosolic subcellular fractions of wildtype and DYT-TOR1A heterozygous MEFs treated with either Veh or 1 μM thapsigargin for six hours. Figure 1 shows a schematic of the workflow used to prepare the experimental samples for proteomic analysis.

**Fig. 1.**
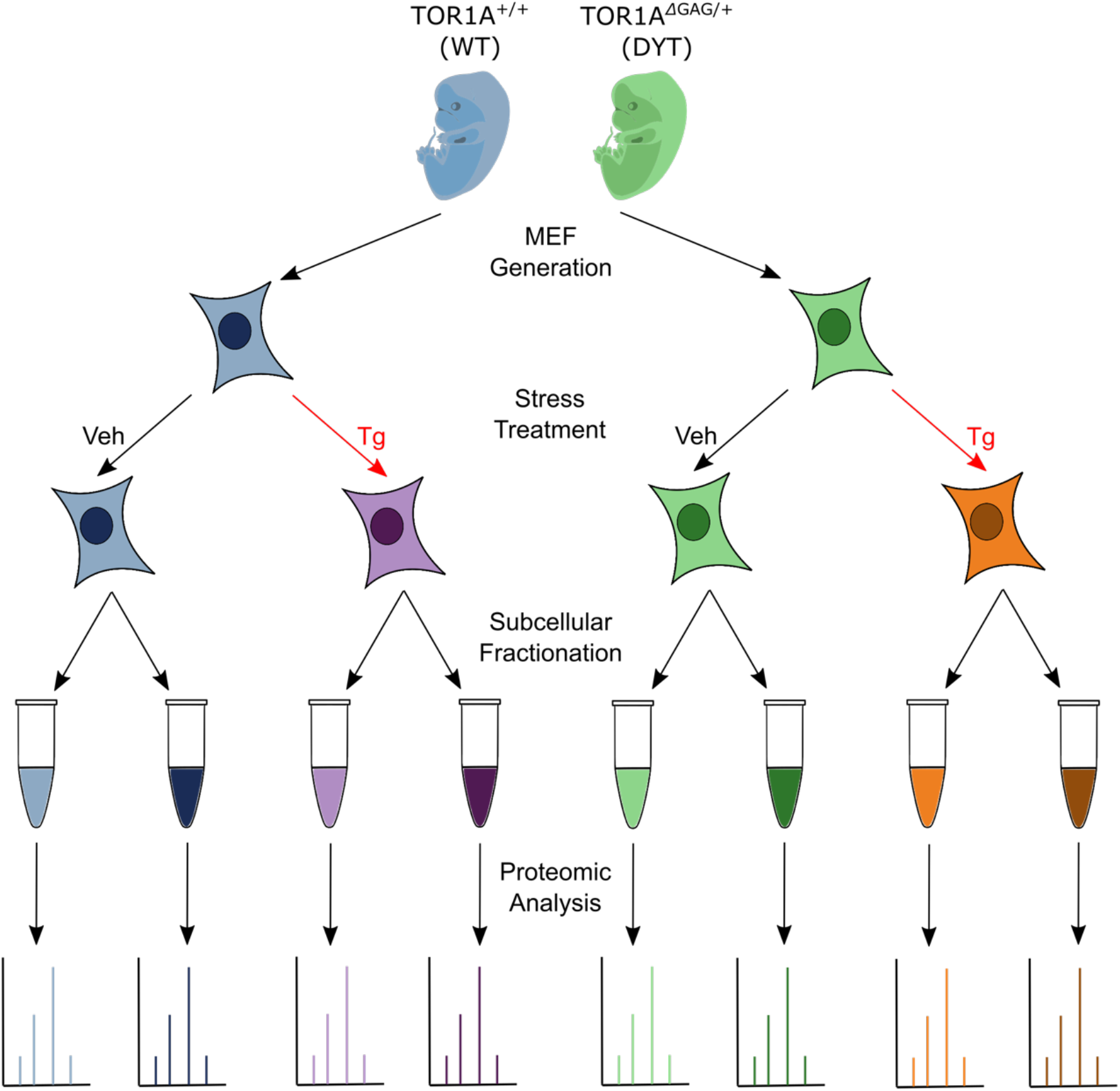
Experimental workflow for proteomic samples. MEFs were generated from heterozygous DYT-TOR1A (*TOR1A*^ΔGAG/+^) mouse embryos and wildtype littermates which were subsequently treated with either 1μM Tg for 6 hours or DMSO vehicle (Veh). Treated MEFs were then fractionated into nuclear and cytosolic fractions and each fraction was subjected to quantitative differential proteomic analysis.

The proteomic profiles for each sample are presented within the supplementary data files. In Supplementary Table 1, we provide the unprocessed mass spectrometry data. In Supplementary Table 2, we provide raw quantitative abundances for all 65,056 unique peptides identified by our proteomics methodology. These abundances are presented for each sample within the experiment. The peptides are grouped within the table based on their alignment with proteins found within the Swissprot *Mus musculus* protein sequence database. In Supplementary Table 3, we present the raw and normalized quantitative abundances for all 4801 unique proteins which had alignments with at least one unique peptide within our peptide dataset. Quantitative abundances for all these proteins are presented for each sample within the experiment, as well as quality control samples. For each protein, separate coefficients of variance are calculated for every experimental condition (i.e. every combination of genotype, treatment, and subcellular fraction). These calculations are based on the normalized quantitative protein abundances from each biological replicate.

## Experimental Design, Materials and Methods

### Animals

DYT-TOR1A knock-in mice *(Tor1a*^ΔGAG/+^)(courtesy of Dr. W. Dauer, University of Texas-Southwestern) [2](Goodchild et al., 2005) on C57BL/6 background were bred in standard housing conditions with food and water provided ad libitum. All procedures were approved by the Duke University Institutional Animal Care and Use Committee (IACUC).

### Mouse Embryonic Fibroblast (MEF) Extraction, Isolation, and Immortalization

Female wildtype C57BL/6 mice were crossed with male heterozygous DYT-TOR1A knock-in mice (*Tor1a*^ΔGAG/+^) on a C57BL/6 background to produce DYT-TOR1A model MEF and WT control MEF cell lines. MEF extraction occurred with minor modifications from the protocol as described in [3](Jozefczuk et al., 2012). Three DYT-TOR1A MEF lines and three WT MEF lines were produced from a single litter using a single animal for each of the different lines.

At approximately 14 days post-coitum, the pregnant dam was euthanized using isoflurane followed by decapitation. The uterine horns were dissected out and rinsed in 70% (v/v) ethanol followed by PBS (Gibco, Invitrogen) before being placed into a Petri dish containing PBS (Gibco, Invitrogen). Each individual embryonic sac was isolated from the uterine horns and placenta, and then placed into a separate Petri dish containing PBS. Within each separate dish, the embryo was dissected out of the embryonic sac.

Following dissection from the embryonic sac, the head and red organs of each embryo were surgically excised. The remaining embryonic tissue was placed into a clean Petri dish where it was minced with a sterile razor blade for 1 minute. 1 mL of 0.05% trypsin/EDTA (Gibco, Invitrogen) was added to each dish containing embryonic tissue. The mixture was transferred into a 15 mL Falcon tube and incubated at 37 °C for 10 minutes. Following incubation, MEFs were dissociated via pipetting. The incubation and dissociation steps were repeated twice more to further dissociate the MEFs from the extracellular matrix of the embryonic tissue. Following the last dissociation step, 2 mL of fetal bovine serum-containing media (“MEF media” described in Cell Culture section below) was added to each tube to inactivate the trypsin. The tubes were subsequently centrifuged at 500 x g for 5 minutes. The supernatant was removed, and the cell pellet was resuspended in 10 mL of warm MEF media.

This solution was then plated on TC dishes coated in 1% Matrigel (Corning). The MEFs were genotyped after two passages and subsequently immortalized via transfection by a plasmid encoding the SV40 T antigen as described in [4](Harding, 2003). Immortalized cell lines were used within 5 passages.

### Genotyping

All genotyping was conducted as previously described in [2](Goodchild et al., 2005).

### Cell Culture

MEFs were grown in MEF media prepared with 500 mL of DMEM, high glucose, pyruvate (Thermo Fisher, #11995), 50 mL of Fetal Bovine Serum, 5 mL of Antibiotic-Antimycotic (Gibco, Invitrogen), 5 mL of 200 mM L-Glutamine (Gibco, Invitrogen), 5 mL of MEM Non-Essential Amino Acids Solution (Gibco, Invitrogen), and 500 μL of 2-Mercaptoethanol (Sigma-Aldrich). MEFs were grown in incubators at 37°C/5% CO_2_.

### Thapsigargin Treatment and Subcellular Fractionation

Three separate experiments were performed exposing MEFs to the cell stressor, thapsigargin (Tg). During each experiment, a pair of MEF lines (1 WT and 1 DYT-TOR1A) were grown within 10 cm dishes to 90% confluence. Treatment consisted of exchanging the MEF media within a dish with MEF media treated with either 1 μM Tg dissolved in DMSO or an equivalent volume of DMSO (Vehicle control, Veh). Following the treatment, the dishes were placed back in incubators at 37°C/5% CO_2_ for six hours. After six hours of treatment, the MEFs were trypsinized to enable dissociation from the 10 cm treatment dishes. The dissociated MEFs were placed in 15 mL Falcon tubes prior to being centrifuged to produce a MEF pellet.

The MEFs were subsequently fractionated into nuclear and cytosolic fractions via a protocol carried out as described in [5](Suzuki et al., 2010). Briefly, the MEF pellet was resuspended in a mild detergent solution consisting of 0.01% NP40 dissolved in PBS. This solution was incubated at room temperature for 3 minutes. Following incubation, the resuspended solution was centrifuged to collect the supernatant (cytosolic fraction). Further solubilization was done in the same buffer alongside sonication to penetrate the double bilayer membranous nuclear compartment. A second centrifugation was performed to collect the supernatant (nuclear fraction).

### Quantitative LC/MS/MS and Proteomic Analysis

Twenty-four samples in total were submitted to the Duke Proteomics and Metabolomics Shared Resource (two subcellular fractions [Nuc, Cyto] from each of the six MEF lines treated with either Tg or Veh). All of the samples were analyzed within a single set of liquid chromatography with tandem mass spectroscopy (LC/MS/MS) data acquistions, irrespective of the treatments occurring as part of separate cell culture experiments.

Fractions were first normalized to 10 μg based on BCA results. Fractions were then spiked with undigested casein at a total of 20, 30, or 40 fmol/μg, reduced with 10 mM dithiothreitol for 30 min at 80 °C, and alkylated with 20 mM iodoacetamide for 30 min at room temperature. Next, they were supplemented with a final concentration of 1.2% phosphoric acid and 741 μL of S-Trap (Protifi) binding buffer (90% MeOH/100mM TEAB). Proteins were trapped on the S-Trap, digested using 20 ng/μL sequencing grade trypsin (Promega) for 1 hour at 47°C, and eluted using 50 mM TEAB, followed by 0.2% FA, and lastly using 50% ACN/0.2% FA. All fractions were then lyophilized to dryness and resuspended in 20 μL 1% TFA/2% acetonitrile containing 12.5 fmol/μL yeast alcohol dehydrogenase (ADH_YEAST). Three QC Pools were created: 1) 3 μL from each of the nuclear fractions (NucPool), 2) 3 uL from each of the cytosolic fractions (CytoPool) 3) 3 μL from each of all of the fractions, both nuclear and cytosolic (TotalPool). All QC Pools were run periodically randomly interspersed throughout the test fractions.

Quantitative LC/MS/MS was performed on 2 μL of each fraction, using a nanoAcquity UPLC system (Waters Corp.) coupled to a Thermo Orbitrap Fusion Lumos high resolution accurate mass tandem mass spectrometer (Thermo) via a nanoelectrospray ionization source. Briefly, the fraction was first trapped on a Symmetry C18 20 mm × 180 μm trapping column (5 μL/min at 99.9/0.1 v/v water/acetonitrile). Analytical separation was then performed using a 1.8 μm Acquity HSS T3 C18 75 μm × 250 mm column (Waters Corp.) with a 90-min linear gradient of 5 to 30% acetonitrile with 0.1% formic acid at a flow rate of 400 nanoliters/minute (nL/min) with a column temperature of 55 °C. Data collection on the Fusion Lumos mass spectrometer was performed in a data-dependent acquisition (DDA) mode of acquisition with a r=120,000 (@ m/z 200) full MS scan from m/z 375 – 1500 with a target AGC value of 2e5 ions.

MS/MS scans were acquired at Rapid scan rate (Ion Trap) with an AGC target of 5e3 ions and a max injection time of 100 milliseconds. The total cycle time for MS and MS/MS scans was 2 seconds. A 20s dynamic exclusion was employed to increase depth of coverage. The total analysis cycle time for each fraction injection was approximately 2 hours.

Following 35 total UPLC-MS/MS analyses (excluding conditioning runs, but including 3 replicate QC Pool (TotalPool), 4 replicate nuclear (NucPool) and 4 replicate cytosolic Pool (CytoPool) injections), data was imported into Proteome Discoverer 2.2 (Thermo Scientific Inc.). Analyses were aligned based on the accurate mass and retention time of detected ions (“features”) using Minora Feature Detector algorithm in Proteome Discoverer. Protein levels are based on the relative peptide abundance measures which were calculated by area-under-the-curve (AUC) of the selected ion chromatograms of the aligned features across all runs. Levels are reported in arbitrary units (a.u.)

Mascot Distiller and Mascot Server (v 2.5, Matrix Sciences) were utilized to produce fragment ion spectra from the MS/MS data and to perform database searches. The MS/MS data was searched against the SwissProt *M. musculus* database (downloaded in Apr 2017) to identify relevant *M. musculus* proteins. The MS/MS data was also searched against an equal number of reversed-sequence “decoys” for false discovery rate determination. Database search parameters included fixed modification on Cys (carbamidomethyl), variable modifications on Meth (oxidation) and Asn and Gln (deamidation), full trypsin enzyme rules, and mass tolerances of 5ppm precursor ions and 0.8da product ions. Peptide Validator and Protein FDR Validator nodes in Proteome Discoverer were used to annotate the data at a maximum 1% protein false discovery rate.

Imputation of missing values was performed in the following manner. If less than half of the values were missing across all samples, values were imputed with an intensity derived from a normal distribution defined by measured values within the same intensity range (20 bins). If greater than half values were missing for a peptide across all samples and a peptide intensity was > 5e6, then it was concluded that peptide was misaligned and its measured intensity was set to 0. All remaining missing values are imputed with the lowest 5% of all detected values.

## Ethics Statement

All animal procedures were approved by the Duke University Institutional Animal Care and Use Committee (IACUC).

## CRediT author statement

KS – Conceptualization, Investigation, Writing – Original Draft, Review & Editing

ZFC – Supervision, Writing – Review & Editing

EJS –Data Curation, Supervision, Writing – Review & Editing

GW – Investigation, Data Curation, Formal Analysis

TH- Investigation, Data Curation

NC – Conceptualization, Funding acquisition, Supervision, Writing – Review & Editing

## Acknowledgments

The authors wish to acknowledge critical expertise and suggestions provided by Shataakshi Dube, William Dauer, Connor King, and Miranda Shipman. K.S. thanks the members of his undergraduate thesis committee, Ron Grunwald and Matthew Oliver, for continued support and guidance.

## Declaration of Competing Interest

The authors declare that they have no known competing financial interests or personal relationships which have or could be perceived to have influenced the work reported in this article.

## Notes

http://dx.doi.org/10.17632/58wghvz7sz.1

### Competing Interest Statement

The authors have declared no competing interest.

