## Supplemental Table 1 for "Dataset on the mass spectrometry-based proteomic profiling of mouse embryonic fibroblasts from a wild type and DYT-TOR1A mouse model of dystonia, basally and during stress"

Due to the size of the .RAW files associated with the unprocessed MS peaks, the data is stored on a separate repository dedicated to holding mass spectrometry data. Please use the information below to access the data.

General Repository: MassIVE

General Repository Link: <https://massive.ucsd.edu/ProteoSAFe/static/massive.jsp>

Dataset Identifier: MSV000087774

Direct Dataset FTP Link: <ftp://massive.ucsd.edu/MSV000087774/>

The FTP link will give you access to the .RAW files containing information about the unprocessed MS peaks from each sample. These samples are labelled using an arbitrary machine ID key (*Ex: ID46169*). Use the key below to associate the machine ID key used in Supplemental Data 1 with the more informative sample label (*Ex: WT_Tg_Nuc_1*) used in Supplemental Tables 2 and 3.

**ID-Label Conversion Key**

*Ex: Table 1 Key = Table 2/3 Label*

ID46169 = WT_Tg_Nuc_1

ID46170 = WT_Veh_Nuc_1

ID46171 = DYT_Tg_Nuc_1

ID46172 = DYT_Veh_Nuc_1

ID46173 = WT_Tg_Nuc_2

ID46174 = WT_Veh_Nuc_2

ID46175 = DYT_Tg_Nuc_2

ID46176 = DYT_Veh_Nuc_2

ID46177 = WT_Tg_Nuc_3

ID46178 = WT_Veh_Nuc_3

ID46179 = DYT_Tg_Nuc_3

ID46180 = DYT_Veh_Nuc_3

ID46181 = WT_Tg_Cyto_1

ID46182 = WT_Veh_Cyto_1

ID46183 = DYT_Tg_Cyto_1

ID46184 = DYT_Veh_Cyto_1

ID46185 = WT_Tg_Cyto_2

ID46186 = WT_Veh_Cyto_2

ID46187 = DYT_Tg_Cyto_2

ID46188 = DYT_Veh_Cyto_2

ID46189 = WT_Tg_Cyto_3

ID46190 = WT_Veh_Cyto_3

ID46191 = DYT_Tg_Cyto_3

ID46192 = DYT_Veh_Cyto_3

ID46193_1 = QC_NucPool_1

ID46193_2 = QC_NucPool_2

ID46193_3 = QC_NucPool_3

ID46193_4 = QC_NucPool_4

ID46194_1 = QC_CytoPool_1

ID46194_2 = QC_CytoPool_2

ID46194_3 = QC_CytoPool_3

ID46194_4 = QC_CytoPool_4
